# SARS-CoV-2 Omicron variant virus isolates are highly sensitive to interferon treatment

**DOI:** 10.1101/2022.01.20.477067

**Authors:** Denisa Bojkova, Tamara Rothenburger, Sandra Ciesek, Mark N. Wass, Martin Michaelis, Jindrich Cinatl

## Abstract

Recently, we have shown that SARS-CoV-2 Omicron virus isolates are less effective at inhibiting the host cell interferon response than Delta viruses. Here, we present further evidence that reduced interferon-antagonising activity explains at least in part why Omicron variant infections are inherently less severe than infections with other SARS-CoV-2 variants. Most importantly, we here also show that Omicron variant viruses display enhanced sensitivity to interferon treatment, which makes interferons promising therapy candidates for Omicron patients, in particular in combination with other antiviral agents.

---

The SARS-CoV-2 Omicron variant (B.1.1.529) causes less severe disease than previous SARS-CoV-2 variants, although immune protection provided by vaccinations and previous infections is reduced against Omicron compared to previous variants [1–4]. In agreement, evidence is emerging that Omicron is inherently less pathogenic than previous SARS-CoV-2 variants. Omicron variant viruses cause less severe disease in animal studies [5–7] and Omicron viruses appear to display a lower capacity than other variants to replicate in the lower respiratory tract [7,8]. Additionally, initial clinical data indicated that the Omicron variant causes less severe disease than previous SARS-CoV-2 variants in unvaccinated individuals [3].

We have most recently shown that Omicron variant viruses are less effective at antagonising the host cell interferon response than Delta variant viruses [9], which provides a mechanistic explanation for the reduced clinical severity of Omicron disease in individuals without pre-existing adaptive immunity [3]. Omicron virus replication was attenuated relative to Delta virus replication in interferon-competent Caco-2 and Calu-3 cells, but not in interferon-deficient Vero cells, and Omicron viruses caused enhanced interferon promotor activity compared to Delta viruses [9]. Additionally, depletion of the pattern recognition receptor MDA5, which plays a critical role in SARS-CoV-2 detection and interferon response initiation [10], resulted in increased Omicron virus replication in interferon-competent cells [9].

The exact molecular reasons for the alleviated interferon response antagonism by Omicron viruses remain to be elucidated. Notably, the Omicron and Delta virus isolates that we investigated (see Suppl. Methods) display sequence variants in the viral interferon antagonists nsp3, nsp12, nsp13, nsp14, the membrane (M) protein, the nucleocapsid protein, and ORF3a [11] (Suppl. Table 1), which may be of relevance.

Here, we show in addition that two SARS-CoV-2 Omicron isolates (Omicron 1, Omicron 2) replicate to lower titres (Figure 1A) and induce elevated STAT1 phosphorylation (Figure 1B), a key event during interferon signalling, compared to a Delta isolate (B.1.167.2) in Caco-2 and Calu-3 cells [9,12] (see Suppl. Methods for further information).

**Figure 1.**
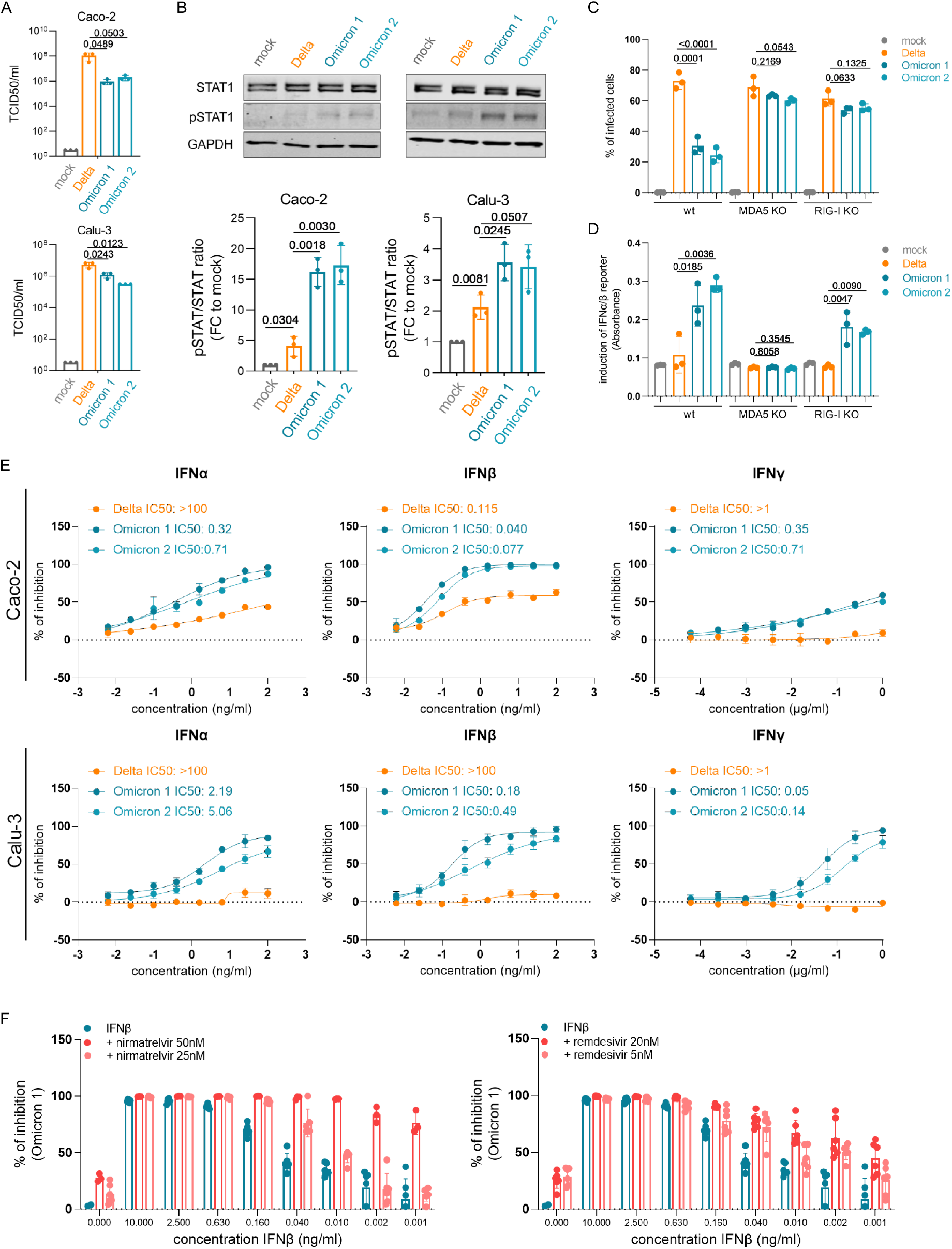
IFN signalling and therapy during infection with novel SARS-CoV-2 variant Omicron. (A) Caco-2 and Calu-3 cells were infected with SARS-CoV-2 variant Delta (GenBank ID: MZ315141), Omicron 1 (GenBank ID: OL800702) and Omicron 2 (GenBank ID: OL800703) at an MOI of 1. The infectious titre was determined 24 h post infection. Graphs represent mean ± SD of three biological replicates. P-values were calculated using Student’s t test. (B) Immunoblot analysis of total and phosphorylated STAT1. The protein levels were quantified by ImageJ. Graphs represent mean ± SD of three biological replicates. P-values were calculated using Student’s t test. (C) A549-ACE2/TMPRSS2 MDA5/RIG-I-WT (wt), A549-ACE2/TMPRSS2 MDA5 KO (MDA5 KO) and A549-ACE2/TMPRSS2 RIG-I KO (RIG-I KO) cells were infected with Delta, Omicron 1 and Omicron 2 variants at an MOI of 0.01 for 72 h. The number of infected cells was determined by immunofluorescence staining. Graphs represent data of four biological replicates. Statistical testing was performed by one-way ANOVA and Dunn’s test. (D) Production of IFNα/β was measured by incubating supernatants from wt, MDA5-KO, and RIG-I KO cells infected with SARS-CoV-2 variants at an MOI of 0.01 for 48 h using HEK-Blue IFNα/β reporter cells. Graphs displays mean ± SD of three biological replicates. Statistical testing was performed by one-way ANOVA and Dunn’s test. (E) Dose response curves of IFNα, IFNβ and IFNγ was assessed in Caco-2 and Calu-3 cells. All IFNs were added to confluent monolayers and subsequently infected with viral variants at MOI 0.01. The inhibition rate was evaluated 24 h (Caco-2) and 48 h (Calu-3) post infection by staining of spike protein. Graphs depicts mean ± SD of three biological replicates. (F) Antiviral effect of IFNβ in combinations with nirmatrelvir and remdesivir in Caco-2 cells.

In A549 cells transduced with ACE2 (SARS-CoV-2 receptor) and TMPRSS2 (mediates SARS-CoV-2 cell entry by cleaving and activating the viral S protein), the Omicron viruses also displayed alleviated infection capacity compared to the Delta virus (Figure 1C). This difference largely disappeared upon depletion of both of the pattern recognition receptors MDA5 and RIG-I that mediate the host cell interferon response in virus-infected cells [13]. However, when we compared interferon activity in the supernatants of SARS-CoV-2-infected cells in a HEK-Blue IFNα/β reporter cell assay, the supernatants of Omicron virus-infected RIG-I-knock out cells induced higher interferon promoter activation than the supernatants of Omicron virus-infected MDA5-knock out cells (Figure 1D). This is in agreement with data showing that MDA5 is primarily responsible for virus recognition and the induction of an interferon response in SARS-CoV-2-infected cells [10,13,14].

Taken together, these findings further confirm that Omicron viruses are less effective than Delta viruses in antagonising the host cell interferon response [9] and that MDA5 is a major player in SARS-CoV-2 recognition [10,13]. Accordingly, elevated MDA5 levels were detected in the upper airways of SARS-CoV-2-infected individuals with mild or asymptomatic disease [15]. Since Delta has been found to display a similar level of interferon antagonism and sensitivity as previous SARS-CoV-2 variants [16,17], the reduced interferon antagonism appears to be unique to Omicron.

Most notably, treatment with interferon-α, interferon-β, and interferon-γ revealed that the weaker interferon antagonism by Omicron virus isolates translates into a profoundly increased Omicron sensitivity to interferon treatment (Figure 1E). Further experiments showed that antiviral interferon-β effects were further increased in combination with nirmatrelvir (the antivirally active agent in paxlovid) and remdesivir (Figure 1F). So far, clinical studies reported mixed outcomes in COVID-19 patients treated with different interferons [18–21]. Given the newly discovered substantially increased interferon sensitivity of Omicron viruses, interferons represent a promising option for the treatment of Omicron patients.

In conclusion, we present further evidence that reduced interferon-antagonising activity explains at least in part why Omicron variant infections are inherently less severe than infections with other SARS-CoV-2 variants. Most importantly, we here also show that Omicron variant viruses display enhanced sensitivity to interferon treatment, which makes interferons promising therapy candidates for Omicron patients, in particular in combination with other antiviral agents.

## Acknowledgements

The authors thank Lena Stegmann, Kerstin Euler and Sebastian Grothe for their technical assistance. This work was supported by the Frankfurter Stiftung für krebskranke Kinder.

## Author contributions

D.B., M.M., and J.C. conceived and designed the study. D.B., T.R., M.N.W., and J.C. performed experiments. All authors analysed data. M.M. wrote the manuscript. D.B., M.N.W., M.M., and J.C. revised the manuscript. All authors have read and approved the final manuscript.

