## Supplemental Materials and Table for "SARS-CoV-2 Omicron variant virus isolates are highly sensitive to interferon treatment"

### Reduced interferon antagonism but similar drug sensitivity in Omicron variant compared to Delta variant SARS-CoV-2 isolates

Denisa Bojkova, Tamara Rothenburger, Sandra Ciesek, Mark N. Wass*, Martin Michaelis*, Jindrich Cinatl jr.*

#### Supplementary information

##### Supplementary methods

###### Cell culture

The Caco-2 (DSMZ, Braunschweig, Germany), Vero (DSMZ, Braunschweig, Germany), Calu-3 (ATCC, Manassas, VA, US) were grown at 37 °C in minimal essential medium (MEM) supplemented with 10% fetal bovine serum (FBS), 100 IU/mL of penicillin, and 100 µg/mL of streptomycin. All culture reagents were purchased from Sigma-Aldrich. A Caco-2 subline of human colon carcinoma cell line that was originally used for the cultivation of SARS-CoV and that is also highly permissive to SARS-CoV-2 [9] was established for isolation and cultivation of SARS-CoV-2 variants as well as for antiviral assays. The A549-ACE2/TMPRSS2 (Invivogen) were grown in DMEM supplemented with 2 mM L-glutamine, 4.5 g/l glucose, 10% (v/v) heat-inactivated fetal bovine serum (FBS; 30 min at 56 ºC), PenStrep (100 U/ml-100 µg/ml), 100 µg/ml Normocin, 10 µg/ml of Blasticidin, 10 µg/ml of Blasticidin, 100 µg/ml of Hygromycin, 0.5 µg/ml of Puromycin, and 100 µg/ml of Zeocin. A549-ACE2/TMPRSS2 MDA5 KO cells (Invivogen) and A549-ACE2/TMPRSS2 RIG-I KO cells (Invivogen) were grown in DMEM supplemented with 2 mM L-glutamine, 4.5 g/l glucose, 10% (v/v) heat-inactivated fetal bovine serum (FBS; 30 min at 56 ºC), PenStrep (100 U/ml-100 µg/ml), 100 µg/ml Normocin, 10 µg/ml of Blasticidin, 100 µg/ml of Hygromycin, 0.5 µg/ml of Puromycin, and 100 µg/ml of Zeocin. All cell lines were regularly authenticated by short tandem repeat (STR) analysis and tested for mycoplasma contamination.

**Virus preparation**

The SARS-CoV-2 isolates Omicron 1 (B.1.1.529: FFM-SIM0550/2021, EPI_ISL_6959871, GenBank ID OL800702), Omicron 2 (B.1.1.529: FFM-ZAF0396/2021, EPI_ISL_6959868, GenBank ID OL800703), and Delta isolate (B.1.167.2: FFM-IND8424/2021, GenBank ID MZ315141) were cultivated in Caco-2 cells as previously described [9,12] and stored at –80°C.

**Determination of infectious titer**

Caco-2 cells and Calu-3 cells were infected with SARS-CoV-2 variants at MOI of 1 for 1h. After the incubation period, the infectious inoculum was removed, cells were washed with PBS and supplemented with fresh medium. One day later, supernatants were collected and stored at -80°C upon titration. To infectious titres were determined by serial dilutions of cell culture supernatants on confluent layers of Caco-2 cells in 96-well plates and expressed as TCID50/ml.

**Immunoblot assay**

Cells were lysed using Triton-X-100 sample buffer (Sigma-Aldrich), and proteins were separated by SDS-PAGE. Detection occurred by using specific antibodies against GAPDH (#2275-PC-100, Trevigen), STAT1 (#9172, CellSignaling), pSTAT1 (#9167, CellSignaling). Protein bands were visualized by laser-induced fluorescence using an infrared scanner for protein quantification (Odyssey, Li-Cor Biosciences, Bad Homburg, Germany). The protein levels were quantified by ImageJ.

**Immunofluorescence staining**

The cells were fixed at indicated times with 3% PFA permeabilized with 0.1 % Triton X-100. Prior to primary antibody labeling, cells were blocked with 5% donkey serum in PBS or 1% BSA and 2% goat serum in PBS for 30 minutes at RT. Spike protein was detected by primary antibody (1:1500, Sinobiological) followed by Alexa Fluor 647 anti-rabbit secondary antibody (1:1000, Invitrogen). The nucleus was labelled using DAPI (1:1000, Thermo Scientific). The images were taken by Spark® Mulitmode microplate reader (TECAN) at 4x magnification.

**Detection of IFNα/β in cell culture supernatants**

A HEK-Blue IFN-α/β (Invivogen) reporter cell line was used to detect the presence of biologically active IFN-α/β in supernatants. Upon IFN-α or IFN-β stimulation, HEK-Blue IFN-α/β cells activate the expression of the reporter gene, SEAP. Briefly, 20 µl of supernatants from SARS-CoV-2 infected cells were incubated together with 180 µl of HEK-Blue IFN-α/β cell suspension (50,000 cells/well) overnight. Then, the expression levels of SEAP were measured by mixing 20 µl of supernatants with 180 µl Quanti-Blue Solution (Invivogen), followed by incubation at 37°C for 5 min and detection at 620 nm by plate reader Infinite 200 (TECAN).

**Immunostaining**

Cells were fixed with acetone:methanol (40:60) solution and immunostaining was performed using a monoclonal antibody directed against the spike protein of SARS-CoV-2 (1:1500, Sinobiological), which was detected with a peroxidaseconjugated anti-rabbit secondary antibody (1:1000, Dianova), followed by addition of AEC substrate. The spike positive area was scanned and quantified by the Bioreader® 7000-F-Z-I microplate reader (Biosys). The results are expressed as percentage of inhibition relative to virus control which received no drug.

**Antiviral assay**

Confluent layers of cells in 96-well plates were treated with decreasing concentration of interferon-α, interferon-β, interferon-γ (all R&D Systems), remdesivir (Selleckchem), and nirmatrelvir (Selleckchem) and subsequently infected with SARS-CoV-2 at an MOI of 0.01. In experiments with remdesivir and nirmatrelvir, 1 µM of the ABCB1 inhibitor Zosuquidar was added. Antiviral effects were determined by immunostaining for the SARS-CoV-2 spike (S) protein 24 h (Caco-2) or 48 h (Calu-3) post infection.

**Statistical analysis**

The results are expressed as the mean ± standard deviation (SD) of number of biological replicates indicated in figure legends. The statistical significance is depicted directly in graphs and the statistical test used for calculation of p values is indicated in figure legends. GraphPad Prism 9 was used to determine IC50 values.

**Supplementary Table 1**. SARS-CoV-2 Omicron-associated sequence variants in proteins described to possess interferon-antagonising activity [11]. Residue numbers are based on the Wuhan reference sequence with residue numbers in brackets indicating the position in the individual proteins within ORF1ab. The residue present in the reference sequence is shown in the reference column. – indicates a deletion.

| **Protein** | **residue** | **Reference** | **OL800702.1** | **OL800703.1** | **Delta** |
|---|---|---|---|---|---|
| ORF1ab (nsp3) | 2083 (1265) | S | - | - | S |
| ORF1ab (nsp3) | 2084 (1266) | L | I | I | L |
| ORF1ab (nsp3) | 2710 (1892) | A | T | T | A |
| ORF 1ab (nsp12) | 4715 (323) | P | L | L | L |
| ORF 1ab (nsp12) | 5063 (679) | G | G | G | S |
| ORF 1ab (nsp13) | 5401 (77) | P | P | P | L |
| ORF 1ab (nsp14) | 5967 (42) | I | V | V | I |
| M | 19 | Q | E | E | Q |
| M | 63 | A | T | T | A |
| M | 82 | I | I | I | T |
| N | 13 | P | L | L | P |
| N | 31 | E | - | - | E |
| N | 32 | R | - | - | R |
| N | 33 | S | - | - | S |
| N | 63 | D | D | D | G |
| N | 203 | R | K | K | M |
| N | 204 | G | R | R | G |
| N | 377 | D | D | D | Y |
| ORF3a | 26 | S | S | S | L |
